# Rapid evolutionary change in trait correlations of a single protein

**DOI:** 10.1101/2022.05.05.490716

**Authors:** Pouria Dasmeh, Jia Zheng, Andreas Wagner

## Abstract

Many organismal traits are genetically determined and covary in evolving populations. The resulting trait correlations can either help or hinder evolvability – the ability to bring forth new and adaptive phenotypes. The evolution of evolvability requires that trait correlations themselves must be able to evolve, but we know little about this ability. To learn more about it, we here study one of the simplest evolvable systems, a gene encoding a single protein, and two traits of this protein, namely the ability to emit yellow and green light. We show that correlations between these two traits can evolve rapidly through both mutation and selection on short evolutionary time scales. In addition, we show that these correlations are driven by a protein’s ability to fold, because single mutations that alter foldability can dramatically change trait correlations. Since foldability is important for most proteins and their traits, mutations affecting protein folding may alter trait correlations mediated by many other proteins. Thus, mutations that affect protein foldability may also help shape the correlations of complex traits that are affected by hundreds of proteins.

## Body

Evolvability is a biological system’s ability to bring forth novel and adaptive phenotypes. Because evolvability varies among organisms and traits, it can itself evolve^1^. Understanding the factors that affect its evolution matters not only for our fundamental understanding of biological evolution. It also matters for technological applications, including the experimental evolution of novel and useful molecules such as efficient industrial enzymes^2^.

Many phenotypic traits are correlated with one another, and these correlations often have a genetic basis^3–5^. Examples of such traits include the weight and the height of individuals within a population^6^. They also include coloration and behavioral traits in many animal species ^3,7,8^, as well as seed dormancy and flower time in plants ^9^. Genetic correlations between phenotypic traits can influence evolvability, because they can affect the extent to which traits can evolve independently from one another^10–14^.

On the one hand, correlated trait evolution can facilitate evolvability. For example, in species of freshwater fish from the genus *Micropterus*, a strong correlation between mouth and jaw morphology helps these species adapt to the consumption of large prey ^15, 16^ On the other hand, strong trait correlations can also hamper evolvability, rendering weak trait correlations advantageous. For example, a weak correlation between early tetrapod forelimb and hindlimb morphology helped bring forth major specializations, such as bipedalism and flight ^17, 18^. More generally, the decoupling of traits from one another helps trait variation become individuated within a population, such that genetic change can affect only one trait without affecting others. Trait individuation can help explain phenomena as different as the evolution of different cell types in multicellular organisms^19^, leaf shapes in plants^20^, and body structures in animals^21^.

Because trait correlations are important for evolvability, it is important to study how easily these correlations can themselves evolve. Previous studies have shown that trait correlations can change on evolutionary time scales, and become weaker or stronger in some lineages of different species. For example, the correlation between beak and the skull morphology in Darwin’s Galapagos finches (genus *Geospiza)* and Hawaiian honeycreepers (*Drepanis coccinea*) is significantly stronger compared to their continental relatives^22^, which has facilitated rapid craniofacial evolution in the former species. Another example is indigenous Australian *dicotyledon* species where the correlations between several seedling traits such as the patterns of scale-like leaves are evolutionarily malleable across thousands of species^23^. Although examples like these show that trait correlations can evolve, we know little about how rapidly they can do so. In addition, we cannot easily quantify the contribution of mutations and selection to these correlations and their evolutionary change. The reason is that they are complex traits of multicellular organisms, which may be influenced by dozens or hundreds of genes. Also, it is difficult to determine the causes of correlations between complex traits. In principle, they can be caused by genetic factors, such as pleiotropy and linkage disequilibrium ^24^, and by environmental and ecological factors, such as habitat temperature and aridity ^25,26^.

To overcome some of these challenges, we study evolutionary change in trait correlations for one of the simplest possible evolving systems, namely a single protein-coding gene. If trait correlations can easily evolve in this simple study system, they would probably be even more malleable in the sets of interacting genes that determine complex macroscopic traits. Our focal gene of interest is *yfp,* which encodes yellow fluorescent protein (YFP). Because this protein plays no important biological role in the microbial host organism *E. coli* in which we study it, we can examine its traits independently from complicating interactions with the genetic background in which it occurs. YFP emits both yellow and green fluorescence light, and these emissions are the two traits we study (Figure 1A). They can be quantified rapidly and precisely in thousands of organisms through fluorescence activated cell sorting (FACS), which also allows us to precisely estimate trait correlations.

**Figure 1.**
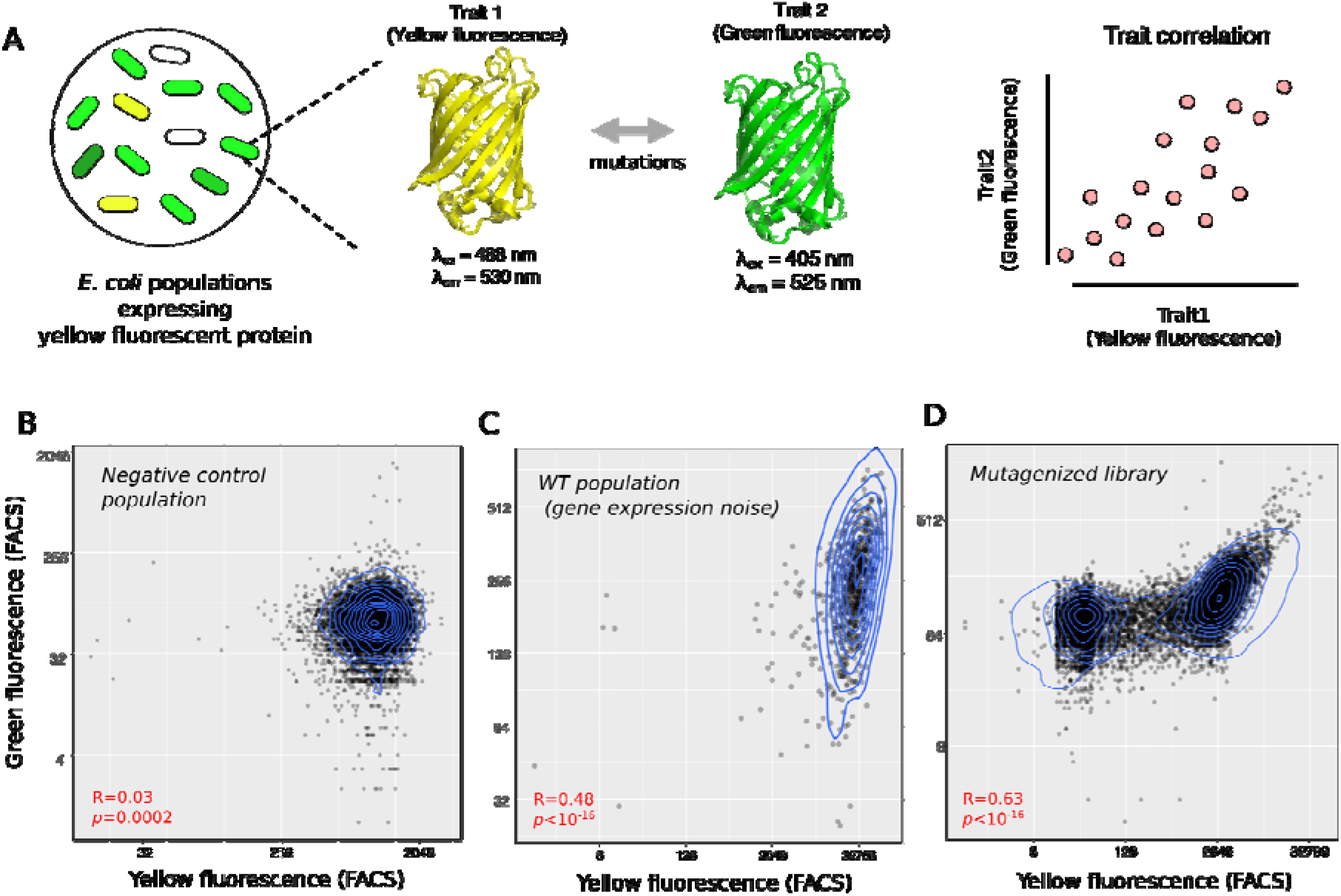
Gene expression noise and mutations can change the strength of correlation between yellow and green fluorescence intensities. A) Schematic of experimental design. We investigate the correlation between yellow and green fluorescence intensities in *E. coli* populations that express yellow fluorescent protein (YFP). B) Autofluorescence correlation between yellow and green fluorescence intensities in an *E.coli* population without the *yfp* gene. C) The correlation between yellow and green fluorescence intensities in an isogenic population expressing wild-type YFP protein. D) The correlation between yellow and green fluorescence intensities in a mutagenized YFP library, where each YFP molecule has, on average, ~ 1 amino acid mutation. Data in panels B-D are based on fluorescence-activated cell sorting (FACS) with ~10^5^ sorted cells (see Methods for details). Blue lines in panels B-D show the contours of a two-dimensional density of data points.

Single point mutations can shift the maximum emission wavelength of YFP. This property facilitates the experimental evolution of fluorescence color in YFP-expressing cell populations. In addition, it allows us to ask whether mutation and/or selection can change the correlation between yellow and green fluorescence. To do so, we systematically measure this correlation in both genetically polymorphic populations of *E.coli* cells that express different YFP variants, as well as in isogenic populations of several YFP mutants. In addition, we also measure this correlation in YFP subpopulations that differ in their fluorescence intensities. These experiments enable us to distinguish the extent to which microenvironmental variation (gene expression noise), mutation, and selection affect the correlation between two traits of a single protein.

Before starting our main experiments, we needed to exclude the contribution of two confounding factors. First, even cells that do not express fluorescent proteins often exhibit some background-fluorescence resulting from other fluorescing molecules ^27^. To exclude the possibility that such autofluorescence may explain the trait correlations we might observe for YFP, we first measured both the yellow and green autofluorescence of *E.coli* cells that do not express YFP. To this end, we used multicolor flow cytometry with two spatially separated lasers, and recorded fluorescence intensity in the yellow and green channels of the cytometer (Methods). The two autofluorescence traits are only weakly correlated (Spearman’s rank correlation R=0.03, p=0.0002;) (Figure 1B). Any significantly stronger correlations in YFP-expressing cells can be attributed to YFP, and not to cellular autofluorescence.

A second confounding factor is a correlation in fluorescence that is not caused by YFP itself but by the environment. Although the external environment (medium and temperature) of our *E.coli* cells did not change during our experiments (see Methods), the microenvironment inside a cell can also affect phenotypic traits. For example, this environment fluctuates constantly due to the thermal molecular motions of its molecules, which causes noisy expression of all genes^28,29^. To identify trait correlations caused by such environmental variation, we measured the correlation between yellow and green fluorescence intensities in a population of isogenic (genetically identical) *E.coli* cells that express wild-type YFP (Figure 1C). Because all cells in such a population have the same genotype, only (micro)environmental variation can contribute to any observed trait correlation in the population. We observed a significant correlation between the two traits (Spearman’s rank correlation R=0.48, *p* < 10^-16^) that was 16 times higher than in autofluorescing cells (R=0.03, *p* < 10^-16^, Spearman’s rank correlation; Figure 1C). This correlation is not unexpected and can be explained by gene expression noise. The reason is that the fluorescence intensity of a single cell is proportional to the abundance of its YFP molecules. Cells with highly expressed YFP will thus fluoresce more intensely in both green and yellow, compared to cells with lowly expressed YFPs, which leads to the substantial environmental trait correlation we observed.

We next turned to one of our central questions, namely whether mutations can strengthen or weaken this baseline correlation. To find out, we measured yellow and green fluorescence in a library of YFP proteins that we had created by error-prone PCR mutagenesis ^30,31^. YFP variants in this library had on average one single point mutation. In this library, we observed a correlation between the two fluorescence intensities (R=0.63, *p* < 10^-16^, Spearman’s rank correlation; Figure 1D), that was significantly stronger than the correlation caused only by gene expression noise (*p* ~ 0, Fisher’s z-transformation). This shows that mutations can readily alter the correlation between our focal traits. We explore the reasons further below.

To investigate whether selection can affect trait correlations as well, we measured the correlation between yellow and green fluorescence intensities in three YFP populations that we had previously evolved under multiple cycles of mutation and either strong selection, weak selection, or no selection for yellow fluorescence intensity ^30,31^ (Figure 2A, see Methods for details). The properties of the YFP protein variants in these populations differ from that of wild-type YFP. For example, the ratio of the median yellow fluorescence intensity of YFPs in populations that evolved under no selection, and either weak or strong selection to that of an isogenic YFP wild-type (WT) population was ~ 0.68, ~ 1, ~18, respectively. Previous single-molecule real-time sequencing had also shown that during this experimental evolution, YFP accumulated up to ~6 amino acid changes compared to the wild-type ^30^.

**Figure 2.**
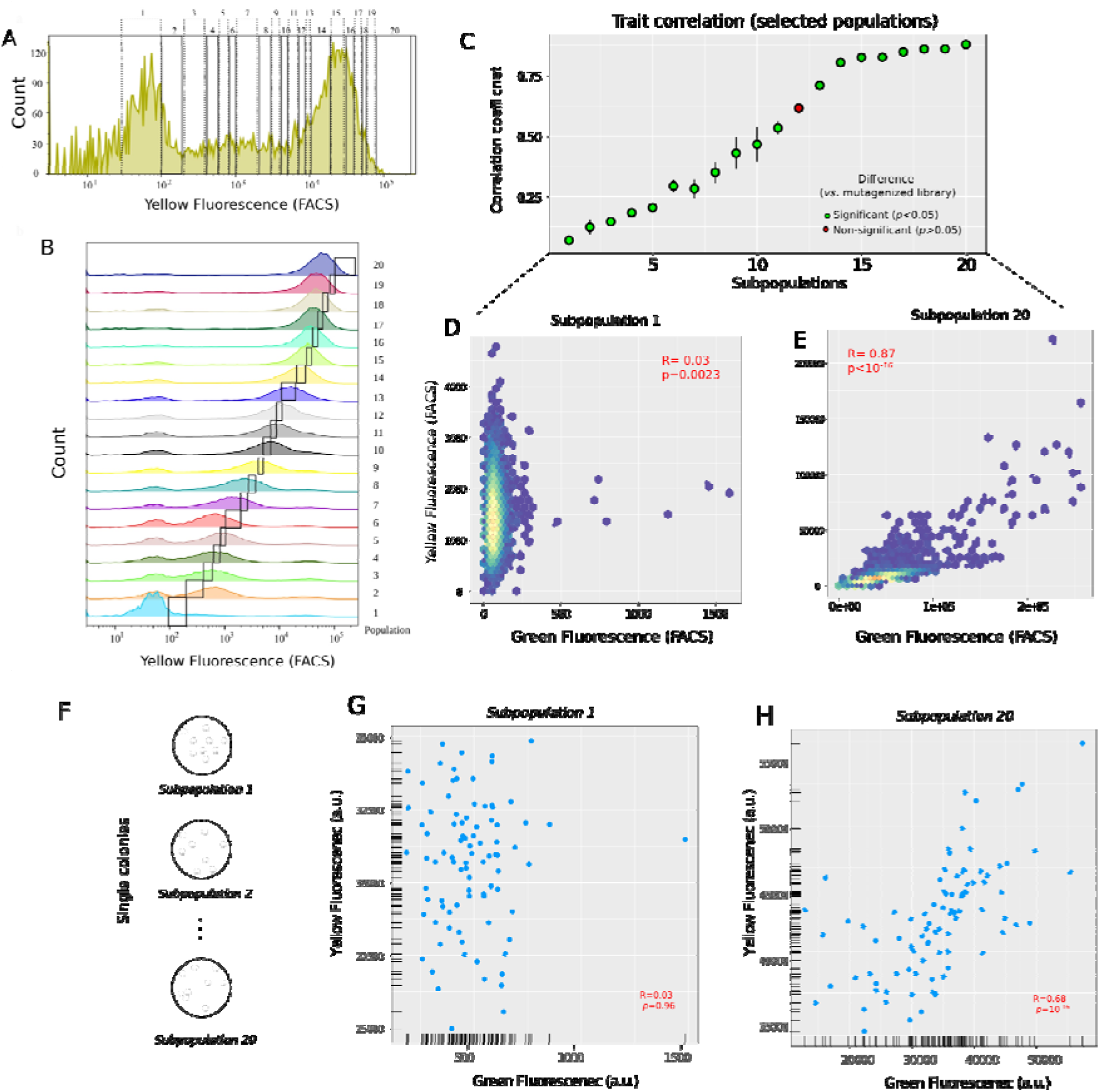
Selection systematically changes the trait correlation measured at the level of single cells. A) Distribution of fluorescence intensity from fluorescence-activated cell sorting (FACS) of our pooled population, which contains YFP variants that have evolved under mutation and various strengths of selection to maintain yellow fluorescence (Methods), and that display different fluorescence intensities. The vertical axis indicates the number of cells at a given yellow fluorescence intensity (horizontal axis, arbitrary units). To study cells of different fluorescence intensity, we sorted cells from this population into twenty bins (subpopulations) according to this yellow fluorescence intensity (Subpopulations 1 to 20, as indicated by lettering on top of each bin). B) Each of the subpanels from bottom to top shows the distribution of fluorescence intensities of cells in one of the 20 subpopulations (right vertical axis). C) Spearman’s correlation coefficient for the correlation between yellow and green fluorescence intensities in all 20 subpopulations (indicated on the x-axis). Error bars represent the standard deviation of the correlation coefficient, as measured from three replicate samples of the same subpopulation. Green circles correspond to subpopulations in which the correlation between yellow and green fluorescence was significantly different from that caused by single point mutations (*p* < 0.05). The red circle corresponds to the 12^th^ subpopulation, in which this correlation was statistically indistinguishable from that caused by mutation (*p* <0.05). We compared the significance of the trait correlation in each subpopulation with the correlation caused by mutations (R=0.63, p < 10^-16^, Spearman’s rank correlation), using Fisher’s z-transformation (see Methods). D) Yellow fluorescence intensity versus green fluorescence intensity for the first subpopulation (with lowest fluorescence intensity). E) Yellow fluorescence intensity versus green fluorescence intensity for the 20^th^ subpopulation (with the highest fluorescence intensity). F) We picked 90 single clones from each of our twenty subpopulations, and measured the correlation between yellow and green fluorescence intensities in liquid cultures derived from each of these 1800 (=90×20) colonies (see Methods). Yellow fluorescence intensity versus green fluorescence intensity for the 90 clones sampled from G) the first subpopulation and H) the 20^th^ subpopulation. All correlation coefficients (R) are Spearman’s rank correlations.

Indeed, trait correlation increased from R=0.5 (*p* < 10^-16^, Spearman’s rank correlation) for the YFP populations that had evolved under no selection, to R=0.78 and R=0.82 (*p* < 10^-16^, Spearman’s rank correlation), for the populations that had evolved under weak and strong selection, respectively (Figure S1). Because we suspected that trait correlations may depend on absolute fluorescence intensities, we pooled YFPs from these three populations and further partitioned this pooled population into 20 subpopulations, such that each subpopulation spanned a similar interval of yellow fluorescence intensity (see Methods). We then sorted ~10^5^ cells in each subpopulation using fluorescence-activated cell sorting (Figures 2B), and measured the correlation between yellow and green fluorescence intensities in each subpopulation.

Remarkably, the correlation between yellow and green fluorescence varied dramatically among the 20 subpopulations. It increased from a weak correlation (R=0.03, *p* = 0.00023; Spearman’s rank correlation) for the first subpopulation with the lowest average yellow fluorescence intensity, to a strong correlation (R=0.87, *p* <10^-16^. Spearman’s rank correlation) for the 20^th^ subpopulation with the highest average yellow fluorescence intensity (Figures 2C-E). The correlation between yellow and green fluorescence was significantly different (*p* < 0.05; Fisher’s z-transformation, Methods) from that caused by mutations (Spearman’s R=0.63) for all except the 12^th^ subpopulation (*p* ~ 0.3, Fisher Z-transformation). These results show that selection can systematically change the strength of a trait correlation, and render it significantly stronger or weaker than the one caused by mutations. To further validate this observation, we picked 90 single clones from each of the 20 subpopulations and measured their yellow (λ_ex_ = 485 nm, λ_em_ = 530 nm) and green (λ_ex_ = 400 nm, λ_em_ = 512 nm) fluorescence intensities using a microplate reader (TECAN Spark) (Figure 2F, see Methods for details). For these single clones too, we observed an increase in the correlation of the green and yellow fluorescence intensities from the first subpopulation to the 20^th^ subpopulation (Figure 2G-H). Altogether, our observations show that trait correlations can be easily shaped by selection, even on the short time scales of laboratory evolution.

We next focused on trait correlations in individual YFP mutants that had appeared in our evolving populations during experimental evolution. We reasoned that these variants might help us find out why trait correlations can vary so strongly from one subpopulation to another. Specifically, we measured trait correlations for 71 YFP mutants that we had previously engineered, because they attained moderate to high frequency in evolving populations ^30,31^. These variants include the WT protein, as well as 10 mutants with one, 28 mutants with two, and 32 mutants with three amino acid changes (Table S1). All double mutations share the mutation G66S (replacement of a glycine with serine at position 66 of YFP) or Y204C, and all triple mutations share both amino acid changes G66S and Y204C. The mutations G66S and Y204C are unique in that they shift the emission spectrum of YFP from yellow towards green fluorescence^30^.

Trait correlations varied significantly among isogenic populations expressing these mutants (Figure 3A). Specifically, they varied from R=0.17 (for the triple mutant G66S-Y204C-F72S; *p* < 10^-16^, Spearman’s rank correlation) to R=0.98 (for the double mutant G66S-N145S, *p* < 10^-16^; Spearman’s rank correlation). Remarkably, some single point mutations sufficed to substantially strengthen (Figure 3B) or weaken (Figure 3C) the trait correlation of the WT protein.

**Figure 3.**
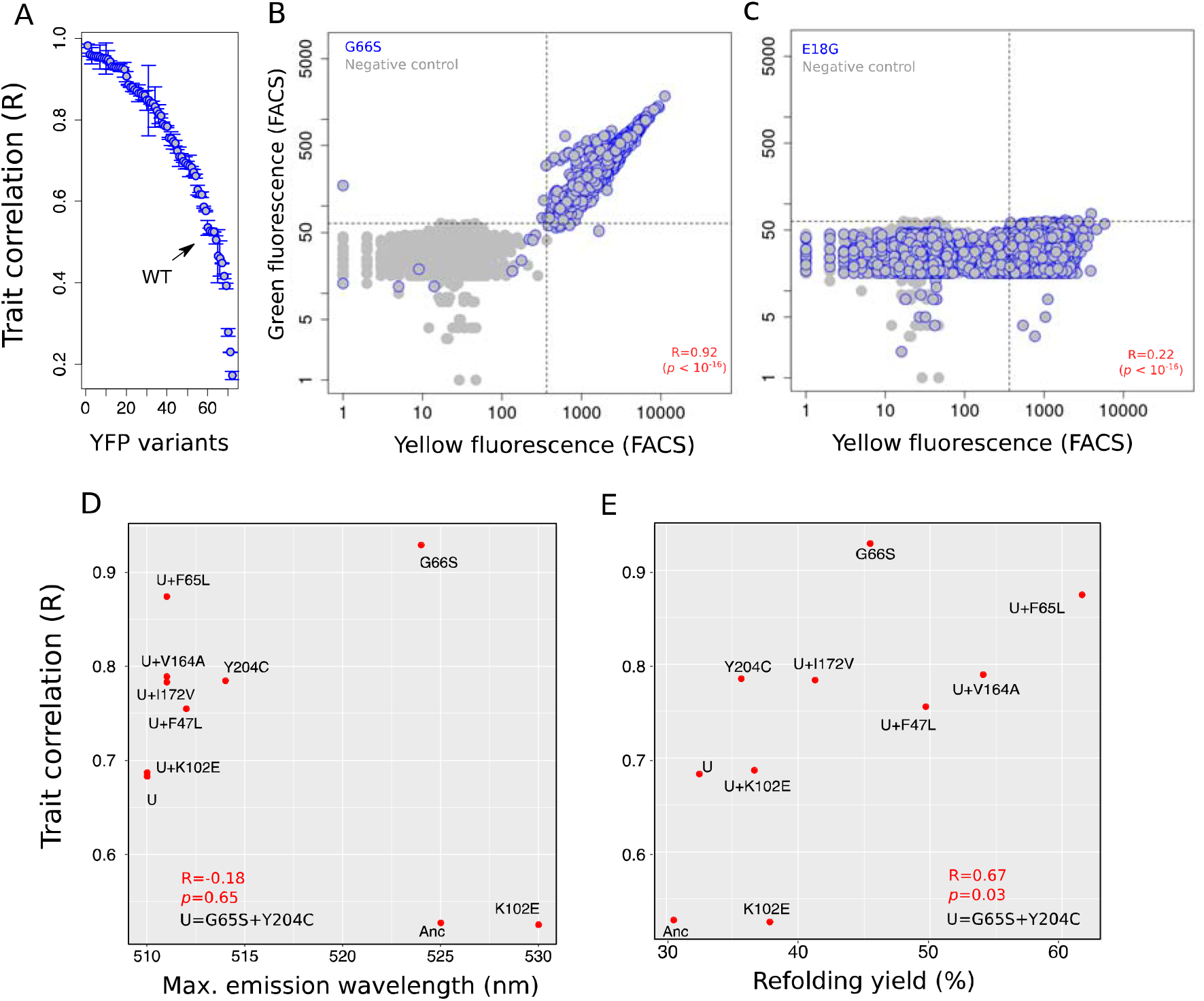
Trait correlation substantially changes by single point mutations that affect protein folding. A) The correlation between yellow and green fluorescence intensities varies substantially in isogenic populations of YFP variants. Each circle corresponds to data from a YFP variant, and error bars show one standard deviation of Spearman’s rank correlation coefficient between yellow and green fluorescence intensities in three biological replicates of isogenic populations expressing the variant.Green fluorescence intensity versus yellow fluorescence intensity for an isogenic population of B) the mutant G66S, and C) the mutant E18G. For both panels B and C, gray circles represent cells from a negative control population that does not express *yfp.* Blue circles represent cells from an isogenic population of the respective mutant. Spearman’s rank correlation coefficient between yellow and green fluorescence intensities versus D) maximum emission wavelength (nm) for a selected set of 10 mutants, and E) refolding yield after thermal denaturation for the same mutants. In panels D-E, the symbol U represents the genotype G66S-Y204C. All correlation coefficients are Spearman’s rank correlations.

We next asked which YFP properties can (i) be easily shaped by one or few mutations, and (ii) affect the trait correlations we observe in individual variants and in our polymorphic subpopulations. One candidate property is fluorescence color itself. Mutations in YFP, particularly those that alter amino acids close to the protein’s fluorophore, can shift the emission spectrum of the protein. We compared trait correlations for a set of 10 YFP variants that maximally emit in colors ranging between yellow (~525-530 nm) and green (~510 nm; see Table S2). However, a change in this emission maximum was not significantly associated with an altered trait correlation (R=0.28, *p*=0.42; Spearman’s rank correlation). For example, although the maximum emission wavelength was substantially different between the mutant G55S (~ 521 nm) and the triple mutants Y204C-G66S-F65L (~ 511 nm), the trait correlation is nearly the same in these two mutants (~ 0.9). Another example involves the triple point mutants that contain both color-shifting mutations Y204C and G66S (~ 511 nm). Among all such mutants, trait correlation varied from 0.68 (for Y204C-G66S-K102E) to 0.87 (Y204C-G66S-F65L). These observations show that the fluorescence color itself. i.e., the emission spectrum, is not the key property affecting the trait correlations we study.

We then focused on a second candidate property, which is biophysical in nature. It is the ability of a protein to fold properly. We hypothesized that changes in protein foldability can affect trait correlations, because only folded proteins fluoresce. To validate this hypothesis, we estimated the foldability of our 10 YFPs variants. Specifically, we measured the overall refolding yield of YFP upon thermal denaturation (see Methods). In this assay, we first denatured YFP, allowed it to refold, and quantified the amount of re-folded proteins by measuring the fluorescence relative to the fluorescence of YFPs that had not denatured^30^. Importantly, more foldable YFP variants showed a significantly higher correlation between green and yellow fluorescence intensities (Figure 3E; R=0.67, *p*=0.03; Spearman’s rank correlation). Some of our 10 YFP variants in this analysis harbored the known foldability-improving mutations F47L, V164A, and F65L^30^. The presence of these mutations alone increased the trait correlation. For example, the trait correlation of the variant G65S-Y20C increased from ~ 0.61 to ~ 0.75, 0.78, and 0.87, in the presence of the mutation F47L, V164A and F65L, respectively (Table S2).

To further validate the hypothesis that changes in foldability determine evolutionary changes in trait correlations, we turned to our 20 subpopulations and quantified the overall refolding yield of YFP upon thermal denaturation for these subpopulations. We also measured the temperature of midpoint denaturation (T_m_) as a measure of thermodynamic stability, another measure that correlates well with protein foldability^32^. Both measures of protein foldability systematically increased from the first to the 20^th^ subpopulation (Figures 4A-B; Figure S2). Both measures were also themselves highly associated with trait correlations (Figures 4C-E; Spearman’s R=0.93 and 0.99 for Tm and refolding yield after thermal denaturation, respectively; *p* ~10^-6^). In addition, we also used an enzyme-linked immunosorbent assay (ELISA) to measure the amount of soluble protein in each subpopulation, which is also a proxy for protein foldability. Indeed, more soluble YFP subpopulations also fluoresced more intensely (Figure S2). But more importantly, protein solubility was again strongly associated with the magnitude of our trait correlations (R=0.99, *p*~10^-6^; Spearman’s rank correlation). Altogether, these results show that protein foldability is a key determinant of the correlation between the two traits we study.

**Figure 4.**
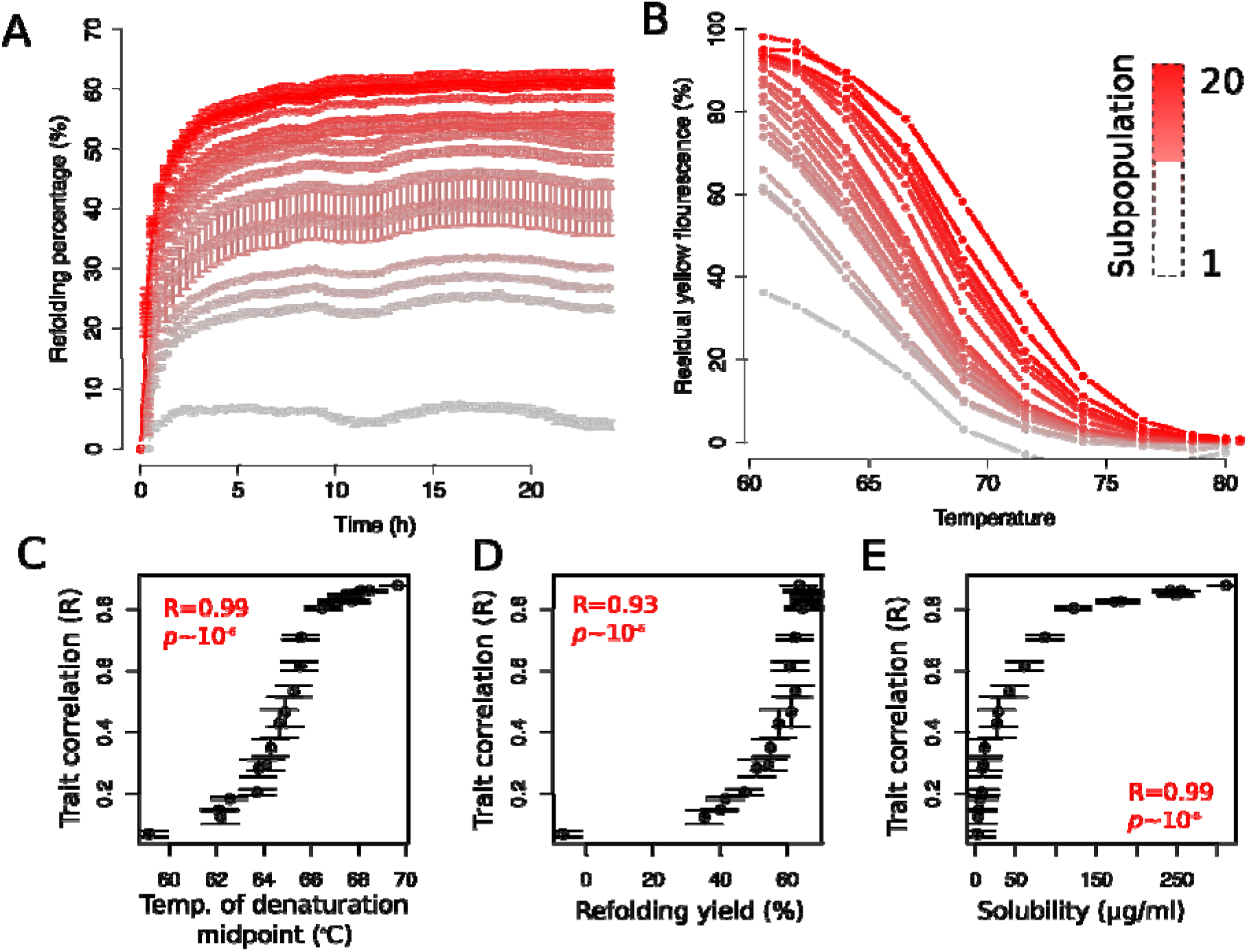
Foldability and solubility of YFP drive trait correlations in YFP populations. A) Refolding percentage upon thermal denaturation for the 20 subpopulations versus time (in hours). B) Residual yellow fluorescence as a function of temperature, for the 20 subpopulations. In panels A and B, subpopulations are color coded from gray (subpopulation 1) to red (subpopulation 20). Spearman’s rank correlation coefficient between yellow and green fluorescence intensities versus C) the temperature of the denaturation midpoint (Tm) of YFP populations, D) the percentage of refolded YFP upon thermal denaturation, and E) the soluble fraction of YFP (in μg/ml) as assessed by an enzyme-linked immunosorbent assay (ELISA). All correlation coefficients are Spearman’s rank correlations.

Taken together, our experiments demonstrate that two traits of a single protein can change dramatically and rapidly on short evolutionary time scales through both mutation and selection. What is more, mutations that alter the stability, foldability, and solubility of proteins can cause this change by affecting the fraction of folded, soluble, and fluorescent proteins in each cell.

A fluorescent protein fluoresces only in its folded state, but like most other proteins, it can exist in a folded, misfolded, or an unfolded state ^32–34^. Any one *E. coli* cell in our experiments contains an ensemble of fluorescent proteins, and within this ensemble, any one protein can randomly transition between these different states. At any instance, each cell thus harbors a fraction of unfolded or misfolded proteins that do not fluoresce. Because these proteins emit neither yellow nor green light, their amount lowers the strength of the correlation between our traits. Indeed, we observed that the fraction of fluorescent cells in our isogenic populations of YFPs not only varied substantially, but was also significantly higher for YFP variants with strong than with weak trait correlations (R=0.87, *p*~10^-6;^ Spearman’s rank correlation; Figure S3). In other words, the more foldable a protein is, the longer it remains in the folded state, and the less likely it is to transit to a misfolded or unfolded states. An ensemble of highly foldable proteins can emit green and yellow fluorescence with fewer fluctuations than an ensemble of less foldable proteins, which leads to a stronger correlation between the traits we study.

If generally true, our observation that single point mutations can readily change a protein’s trait correlations also has implications for correlations between complex traits of higher organisms, and their ability to change over longer evolutionary time scales. Such traits are affected by multiple proteins, and if single mutations can alter trait correlations mediated by at least some of these proteins, these correlations may also be highly malleable in evolution. A recent study that compared correlations between morphological traits of the budding yeast *Saccharomyces cerevisiae* supports the malleability of xomplex traits. It compared correlations in morphological traits observed in single-gene deletion lines of yeast with correlations between the same traits observed in 16 different yeast strains^35^. Many trait pairs showed significantly different correlations in the deletion mutants than in the yeast strains, which shows that the correlations between complex traits that involve multiple genes can readily change on the time scales on which new yeast strains evolve.

The main biophysical property that make trait correlations malleable in our experiments is foldability. This property is essential for the function of most proteins^36–39^. It can also readily change by mutations. Specifically, most random mutations in proteins are destabilizing^40^, but ~20% of such mutations are stabilizing and increase protein foldability^40^. In consequence, protein foldability is highly variable during protein evolution^41–44^. For example, even though the ability to bind oxygen does not differ substantially among mammalian myoglobins (oxygenation constant ~ 0.8-1.2 μM^-1^), the unfolding transitions of these myoglobins differ nearly 600-fold in their resistance to chemical denaturants^44,45^. Other well-studied proteins include the essential bacterial dihydrofolate reductase (DHFR), which reduces dihydrofolic acid to tetrahydrofolic during folate biosynthesis, as well as mammalian superoxide dismutase 1 (SOD1), which reduces superoxides to hydrogen peroxides to destroy damaging free superoxide radicals. Mutations that affect the stability of these proteins’ secondary and tertiary structures also alter their foldability ^46–48^. More generally, proteins in the proteomes of *E. coli, C. elegans, S. cerevisiae,* and human vary widely in their thermodynamic stability^49,50^, which correlates with protein foldability^32^. These examples show that changes in protein foldability are frequent in the evolution of proteins. As our work showed, such changes can also lead to substantial changes in trait correlations. Such malleable trait correlations caused by changes in protein foldability can facilitate the co-evolution or individuation of simple or complex traits in evolution.

In sum, our experiments not only show that correlations between traits of a single protein can change rapidly and on short evolutionary time scales. They also provide a simple biophysical explanation for this change. They link the fundamental protein property of foldability with a fundamental aspect of evolvability – trait correlations. In doing so, they can help explain how evolvability can evolve rapidly.

## Data availability

Scripts, raw and processed data, and statistical analyses are available at:https://github.com/dasmeh/yfp_trait_correlation

## Funding and competing interest

This project has received funding from the European Research Council under Grant Agreement No. 739874. We would also like to acknowledge support by Swiss National Science Foundation grant 31003A_172887 and by the University Priority Research Program in Evolutionary Biology at the University of Zurich.

## Materials and Methods

### Plasmids, strains, and mutant libraries

We used the vector pBAD202/D-TOPO (K4202-01, Invitrogen), which carries an arabinose-inducible araBAD promoter and a Kanamycin-resistance gene for YFP evolution within the E. coli strain BW27783 (CGSC 12119). As described previously^3^, we inserted the coding region of *yfp* from water jellyfish *(Aequorea victoria;* Uniprot ID: A0A059PIR9) that was already present in the plasmid pAND^2^ into the vector backbone of pBAD202/D-TOPO by restriction digest and ligation. We inserted the YFP-coding gene between *XhoI* and *HindIII* restriction sites, placing it under the control of the arabinose-inducible *araBAD* promoter. We named the resulting plasmid pBAD-YFP.

We introduced random mutations into the coding region of YFP by mutagenic PCR as previously described^3^. Briefly, we added 10 ng of template plasmid to 100 μl of a PCR reaction mix that contains 10 μl of 10×ThermoPol buffer (M0267L, NEB), 2.5 μl of *Taq* DNA polymerase (M0267L, NEB), 400 μM of dNTPs (R0192, Thermo Scientific), 3 μM of 8-oxo-GTP/dPTP (Trilink Biotechnologies), and 400 nM of each primer (MutafpF-GAAGGAGATATACctcgag /MutafpR- AGACCGTTTAAACaagctt). We used a Biometra thermocycler to perform PCR by using the following program: 95 °C/30 s; 25 cycles of 94 °C/20 s, 46 °C/30 s and 68 °C/50 s; 68 °C/5 min. We used the restriction enzymes *Xho*I and *HindIII* (R0146L/R3104S, NEB) to digest the resulting PCR products, and used *DpnI* (R0176S, NEB) to remove the template plasmid by following the manufacturer’s protocols. We used the QIAquick PCR purification kit to purify the digested mutated YFP pools to obtain linearized inserts.

### Sorting cells from a pool of evolving populations at the end of directed evolution

To systematically study how selection affects protein evolvability through direct effects on fitness and indirect effects on protein stability and foldability, we sampled diverse YFP variants that vary broadly in yellow fluorescence. Specifically, we sampled from YFP populations in a previously published directed evolution experiment, in which we had evolved populations of YFP under either strong selection for yellow fluorescence (populations S, top 20 percent of yellow fluorescing cells survive selection), weak selection (populations W, top 65 percent survive), or no selection (populations N for neutral, 100 percent survive)^30^. For each of these selection regimes we had evolved four replicate populations for four rounds (“generations”) of directed evolution through fluorescent activated cell sorting (FACS)-based selection and PCR-based mutagenesis.

To create a pool of yellow fluorescent proteins that cover a broad range of fluorescence intensities, we sampled 100 μl of glycerol stock from each of the four replicate S populations (from generation 4), W populations (generation 4), and W populations (generation 2). We added to each of these 12 (=4+4+4) samples 2 ml LB medium supplemented with 30 μg/ml kanamycin. We grew the resulting 12 cultures at 37 °C with shaking at 220 rpm in a 10 ml tube for ~5 h, and then transferred 50 μl of each culture to 2 ml LB supplemented with 30 μg/ml kanamycin. After continuing cultivation for another ~12 h, we combined 400 μl of each culture into one tube and mixed thoroughly. Subsequently, we added 900 μl of the resulting mixture into 600 μl 50% glycerol and stored it at −80 °C for the subsequent sorting experiments. We call the resulting mixture our “pooled” population of YFP-expressing cells. We sorted this pooled populations into 20 “subpopulations” as shown in Figure 2A.

To sort the pooled population into 20 subpopulations according to their fluorescence, we first added 200 μl of glycerol stock of the pooled population sample to 3 ml LB medium containing 30 μg/ml kanamycin, and grew the resulting culture at 37 °C with shaking at 220 rpm for ~5 h. We then transferred 200 μl of the culture into 20 ml LB medium with 50 μg/ml kanamycin, and continued the incubation for ~12 h. We then sampled 2 ml of the culture and centrifuged it at 9,000 g and 4 °C for 5 min to collect cells. We suspended the collected cells in 2 ml LB medium supplemented with 50 μg/ml of kanamycin and 0.2% arabinose, and continued cultivation for ~12 h. Subsequently, we sampled 40 μl of culture and suspended it in 2 ml cold PBS buffer. We selected cells by their yellow fluorescence intensity according to the selection criteria described in Figures 2A-B with an Aria III cell sorter (BD Biosciences). Specifically, we sorted cells at 4 °C in the FITC channel (λex = 488 nm, λem = 530±15 nm), and collected 10^5^ cells in ~1 ml LB medium for each sorted subpopulation. We placed the selected cells on ice until we had finished sorting all subpopulations to prevent cell proliferation or death. We prepared a glycerol stock of each subpopulation for later flow cytometry measurements.

### Fluorescence assay using flow cytometry

We added 200 μl of glycerol stock from each subpopulation into 10 ml LB medium containing 50 μg/ml of kanamycin, and incubated the resulting 20 cultures at 37 °C with shaking at 220 rpm for ~8 h. We then sampled 2 ml of each subpopulation’s culture, and centrifuged it at 9000 g and 4°C to collect cells. To the pelleted cells of each subpopulation we added 2.3 ml LB medium supplemented with 50 μg/ml of kanamycin and 0.2% arabinose, resuspended the cells, and grew the resulting culture at 37 °C with shaking at 220 rpm for ~12 h. This procedure yielded 20 overnight cultures. We then added 20 μl of these 20 overnight cultures to 180 μl of cold PBS buffer. After mixing thoroughly by pipetting, we transferred 5 μl of the resulting suspension into 195 μl of cold PBS buffer, mixed thoroughly, and measured yellow fluorescence in the FITC channel (λ_ex_ = 488 nm and λ_em_ = 530 ± 15 nm) and green fluorescence in the AmCyan channel (λ_ex_ = 405 nm and λ_em_ = 525 ± 25 nm) at room temperature. We used a Fortessa cell analyzer (BD Biosciences) to analyze ~10^4^ events per biological replicate with a flow rate of ~3000 events/s. We sorted three biological replicates for each subpopulation We placed all samples on ice during sample analysis to prevent cell proliferation or death.

### Flow cytometry data analysis

We used FlowJo V10.4.2 (LLC) to analyze flow cytometry data. Specifically, we first selected a homogenous cell population by forward scatter height (FSC-H) versus side scatter height (SSC-H) density plots. We then selected singlets (single cells) by using side scatter area (SSC-A) versus side scatter height (SSC-H) density plots. We used the resulting filtered data for determining green and yellow fluorescence intensities.

### Extracting soluble fluorescent proteins

After inducing the expression of YFP variants in each subpopulation as described in *Fluorescence assay using flow cytometry,* we sampled 2 ml of the overnight culture of each subpopulation for extracting soluble fluorescent proteins by following the manufacturer’s protocol. Specifically, we centrifuged the culture for each subpopulation at 5,000g and at 4 °C for 5 min to collect cells, and stored the collected cells at −20 °C overnight. We then followed the manufacturer’s protocol to extract soluble proteins by using CelLytic™ B Cell Lysis Reagent (B7435-500ml, Sigma). Subsequently, we used 200 μl cell lysis solution (ThermoFisher; 50 mM Tri with pH 7.4, 250 mM NaCl, 5mM, 50 mM NaF, 1mM Na_3_VO_4_, 0.02% NaN3) to dissolve the soluble proteins, and placed the resulting soluble pellet sample on ice for subsequent experiments.

### Protein refolding assay

To unfold fluorescent proteins, we mixed 5 μl of crude lysate of each subpopulation with 45 μl of 8M urea (containing 10 mM DTT), and heated the sample at 95 °C for 5 min in a PCR thermocycler. Asa control, we mixed 5 μl of crude lysate with 45 μl of TNG buffer (100 mM Tris, 100 mM NaCl, 10% glycerol, 10 mM DTT, 1×cOmplete™ (EDTA-free Protease Inhibitor Cocktail, Roche 11873580001), pH 7.2-7.5). To refold the unfolded fluorescent proteins, we rapidly added 10 μl aliquots of an unfolded sample or of the control into 190 μl of TNG buffer in a 96 well microplate, and immediately measured fluorescence intensity using an infinite F200 Pro microplate reader (λex = 485 nm, λem = 530 nm). We measured fluorescence at ~20 min intervals with 2-mm orbital shaking in between. We report the refolding yields as fluorescence relative to the control.

### Protein thermal stability assay

To quantify the thermal stability of fluorescent proteins in each subpopulation, we added 2 μl of crude lysate to 98 μl of TNG buffer, and mixed thoroughly by pipetting. We incubated the resulting mixture in a PCR cycler for 5 min, and subjected each subpopulation to a temperature range of 60.6-80.6 °C (specifically, 60.6, 62, 64.1, 66.6, 69, 71.6, 74, 76.5, 78.6, 80 and 80.6 °C), followed by a 30 second incubation at 4 °C. Then we immediately transferred 90 μl of each mixture to a 96 well microplate, and used an infinite F200 Pro (λex = 485 nm, λem = 530 nm) to measure its fluorescence intensity. As a control, we used the unheated lysate-buffer mixture. We report thermal stability as fluorescence relative to the control.

### Quantification of soluble fluorescent proteins by ELISA

To quantify the amount of soluble fluorescent proteins in each subpopulation, we used a GFP ELISA Kit (AKR-121, Cell Biolabs Inc.) which can detect GFP, BFP, CFP, and YFP from *Aequorea victoria.* Specifically, we followed the manufacture’s protocol to determine the quantity of soluble fluorescent proteins in the lysate of each biological replicate for every subpopulation by comparing its absorbance with that of a recombinant GFP standard curve.

### Fluorescence assay using a microplate reader

To further validate how different selection strengths affect the association between green and yellow fluorescence, we randomly sampled 90 clones from each of the twenty subpopulations, and measured their green and yellow fluorescence intensities using a microplate reader. Specifically, we used saline to dilute the glycerol stock of each subpopulation 10^5^-fold, and plated 100 μl of the resulting culture on LB agar supplemented with 25 μg/ml kanamycin. We incubated the LB agar plates in an incubator at 37 °C overnight, picked single colonies, and inoculated each colony into 200 μl of LB medium (50 μg/ml kanamycin). We grew the resulting cultures in a microplate incubator at 37 °C and 1000 rpm. After ~5 h of incubation, we transferred 50 μl of each culture into 150 μl LB medium supplemented with 0.2% (w/v) arabinose and 50 μg/ml kanamycin, and continued the incubation for ~16 h. We then mixed 50 μl of each culture with 170 μl PBS buffer by pipetting thoroughly, and used a TECAN microplate reader (TECAN Spark) to measure both yellow (λex = 485 nm, λem = 530 nm) and green (λex = 400 nm, λem = 512 nm) fluorescence intensities.

### Statistical analyses

To test the null hypothesis that trait correlations are identical between samples (e.g., when comparing trait correlations between two subpopulations), we used Fisher’s z-transformation. In this approach, Pearson’s or Spearman’s correlation coefficients are converted to z-scores, so that they become normally distributed. The null hypothesis is then tested using a t-test on the z-scores. We performed all statistical analyses with R (v.4.0)^51^.

## Notes

### Competing Interest Statement

The authors have declared no competing interest.

