## Supplementary Information for "Rapid evolutionary change in trait correlations of a single protein"

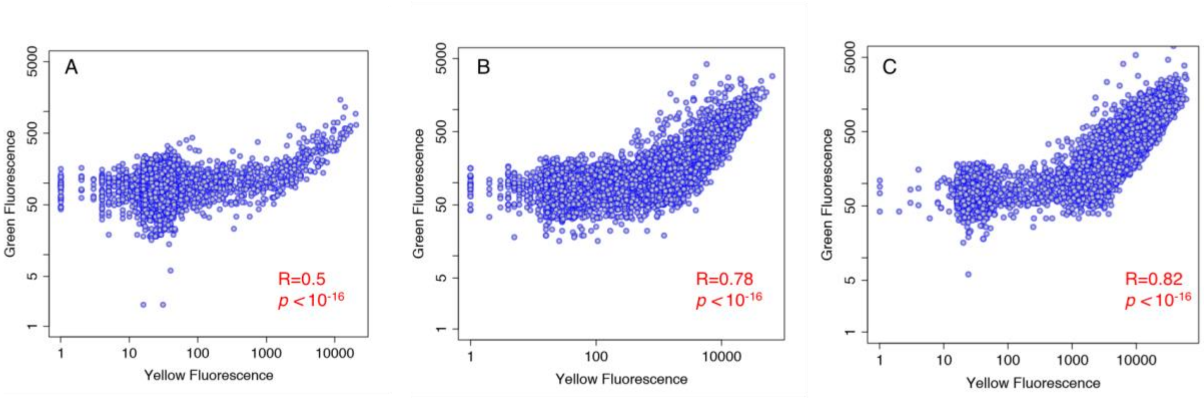


**Figure S1.** Green versus yellow fluorescence intensities for polymorphic populations that had evolved under A) no selection for yellow selection, B) weak selection (top 65 percent survive), and C) strong selection (top 20 percent survive). All correlation coefficients are Spearman's rank correlations.

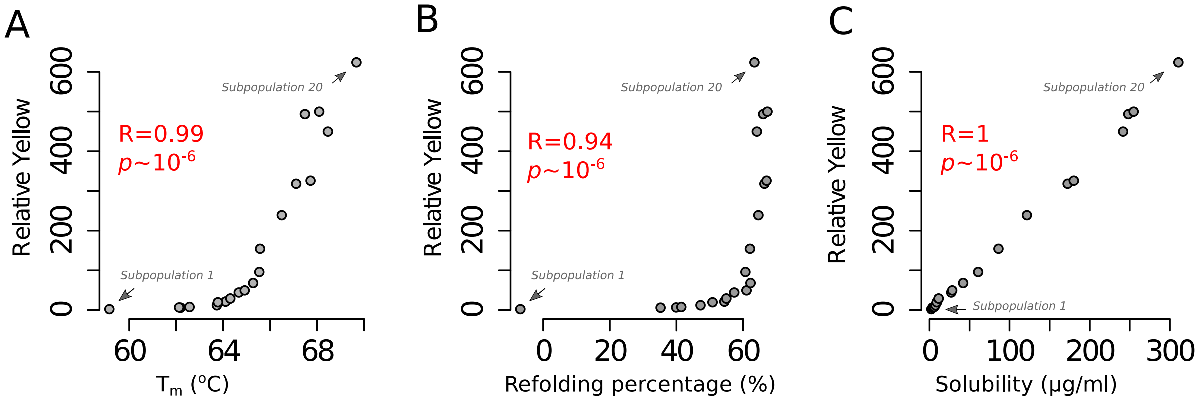


**Figure S2.** Yellow fluorescence intensity (relative to a negative control population) is plotted for all subpopulations (vertical axis) versus A) the temperature of the denaturation midpoint (Tm) of YFP from each subpopulation, B) the refolding percentage upon thermal denaturation, and C) The soluble fraction of YFP proteins (in µg/ml), as assessed by an enzyme-linked immunosorbent assay (ELISA). In panels A-C, subpopulation number increases from the first subpopulation (the left-most circle, labeled), to the 20th subpopulation (right-most circle, labeled). All correlation coefficients are Spearman's rank correlations.

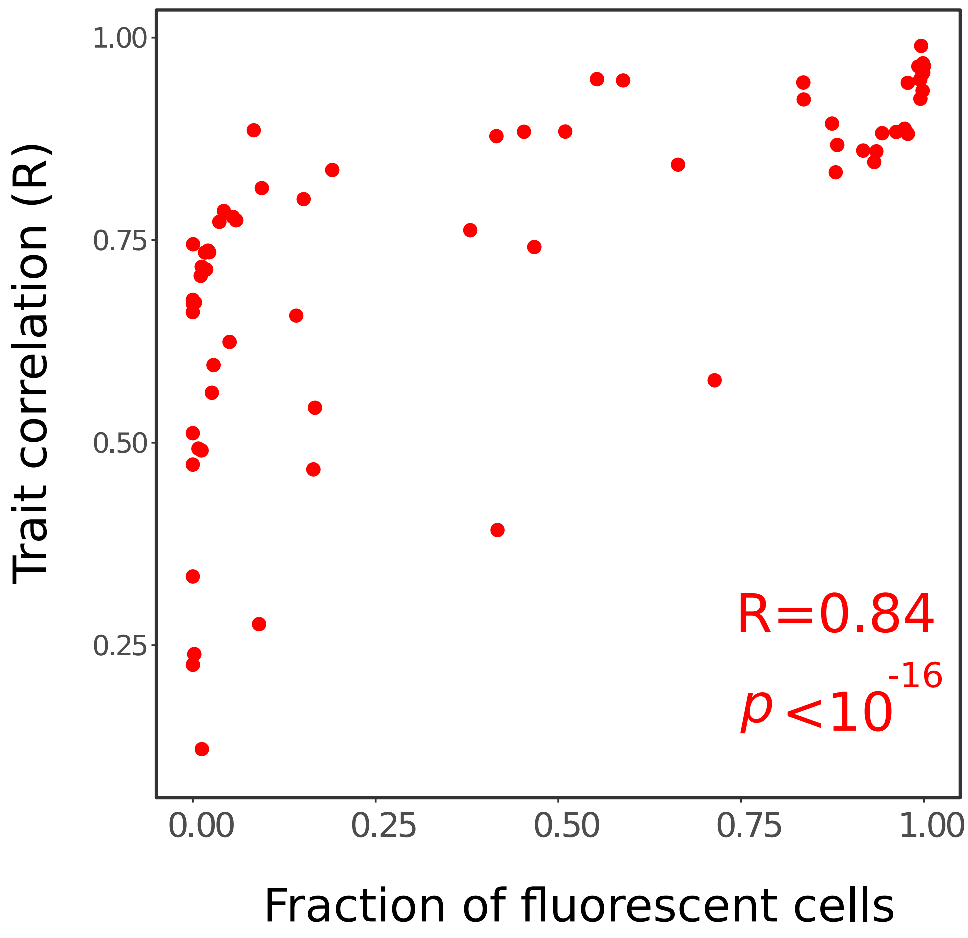


**Figure S3.** The correlation between yellow and green fluorescence intensities versus the fraction of fluorescent cells for 71 YFP mutants that we had previously engineered, because they attained moderate to high frequency in evolving populations 1,2. These variants include the WT protein, as well as 10 mutants with one, 28 mutants with two, and 32 mutants with three amino acid changes (Table S1). All double mutations share the mutation G66S (replacement of a glycine with serine at position 66 of YFP) or Y204C, and all triple mutations share both amino acid changes G66S and Y204C. We calculated the fraction of fluorescent cells by counting the number of cells whose fluorescence was higher than the maximum fluorescence intensity of cells within our negative control population. The correlation coefficient is Spearman's rank correlations.

**Table S1.** Spearman's rank correlation coefficient R between green and yellow fluorescence intensities of YFP mutants. The symbols G, Y and U correspond to the mutations G66S, Y204C, and G66S-Y204C, respectively.

| Mutation | R[a] | Mutation | R | Mutation | R |
|---|---|---|---|---|---|
| U_F72S | 0.172 | U_K141R | 0.698 | U_F65L | 0.874 |
| E18G | 0.230 | U_I129T | 0.709 | U_V2A | 0.880 |
| U_M79L | 0.278 | U_K167E | 0.709 | U_V2A | 0.880 |
| U_E18G | 0.392 | U_G5S | 0.723 | G_F65S | 0.886 |
| M79L | 0.416 | U_R74H | 0.742 | U_I168V | 0.906 |
| R110S | 0.448 | U_F65S | 0.746 | Y_V2M | 0.923 |
| U_R110S | 0.460 | U_K102R | 0.753 | Y_V2A | 0.925 |
| G5S | 0.465 | U_F47L | 0.755 | U_V2M | 0.927 |
| E7K | 0.505 | U_I172V | 0.783 | U_V2M | 0.927 |
| U_F65S | 0.525 | Y204C | 0.784 | G66S | 0.929 |
| K102E | 0.525 | U_V164A | 0.789 | G_F47L | 0.929 |
| WT | 0.527 | Y_N145S | 0.809 | G_F72I | 0.930 |
| K157E | 0.535 | Y_F72I | 0.814 | G_F72C | 0.943 |
| G233D | 0.576 | U_T60S | 0.822 | G_K141R | 0.950 |
| U_K157E | 0.586 | Y_V164A | 0.832 | G_V164A | 0.951 |
| U_G233D | 0.616 | Y_F47L | 0.840 | G_K102R | 0.951 |
| U_K42R | 0.617 | Y_K141R | 0.842 | G_V2A | 0.954 |
| U_E7K | 0.628 | Y_K102R | 0.847 | G_I172V | 0.955 |
| U_E91A | 0.662 | Y_I129T | 0.848 | G_F65L | 0.956 |
| U_I48V | 0.671 | Y_F72C | 0.859 | G_I168V | 0.956 |
| U | 0.683 | U_F72I | 0.861 | G_I129T | 0.958 |
| U_K102E | 0.687 | Y_I172V | 0.864 | G_V2M | 0.960 |
| U_N145S | 0.689 | U_F72C | 0.865 | G_N145S | 0.982 |
| Y_I168V | 0.694 | Y_F65L | 0.871 | | |

**Table S2.** Maximum emission wavelength (nm), refolding yield after thermal denaturation (%), and Spearman's rank correlation coefficient R between green and yellow fluorescence intensities of a selected set of 10 YFP mutants.

| Mutant | Max. emission wavelength (nm) | Refolding yield (%) | Trait correlation |
|---|---|---|---|
| G66S | 524 | 45.45 | 0.9285 |
| WT | 525 | 30.56 | 0.5271 |
| Y204C | 514 | 35.68 | 0.7844 |
| K102E | 530 | 37.84 | 0.5252 |
| G66S-Y204C | 510 | 32.5 | 0.6828 |
| G66S-Y204C -F47L | 512 | 49.66 | 0.7546 |
| G66S-Y204C -F65L | 511 | 61.5 | 0.8738 |
| G66S-Y204C -V164A | 511 | 54.01 | 0.7888 |
| G66S-Y204C -I172V | 511 | 41.27 | 0.7829 |
| G66S-Y204C -K102E | 510 | 36.67 | 0.6868 |
